# A Chitin-Binding Protein Ultra-highly expressed in the Outer Fold of Mantle is Related to Shell Colour in Pacific Oyster *Crassostrea gigas*

**DOI:** 10.1101/2022.04.18.488701

**Authors:** Mai Li, Juyan Tang, Baoyu Huang, Yaqiong Liu, Lei Wei, Yijing Han, Xiaona Wang, Xiuxiu Sang, Mengqiang Yuan, Nini Fan, Shuai Cai, Yanxin Zheng, Meiwei Zhang, Xiaotong Wang

**Affiliations:** School of Agriculture, Ludong University, Yantai, China; Changdao Enhancement and Experiment Station, Chinese Academy of Fishery Sciences, Yantai, China

**Author notes:** Correspondence: Meiwei Zhang, Xiaotong Wang. These authors have contributed equally to this work (Mai Li, Juyan Tang).

## Abstract

Molluscs constitute the second largest phylum of animals in the world, and shell colour and stripes are one of their most important phenotypic characteristics. Studies on the mechanism of shell pigmentation help understand the evolutionary and ecological significance of shell colour and serve as the basis for shell colour breeding. In this study, a matched-pair design was used in comparing the black- and white-striped mantles of the same oyster. The result showed that the stripes of shell surface are corresponding to the stripes of the mantle edge. Transcriptomic analysis identified an uncharacterized protein gene (we named it as *Crassostrea gigas* chitin-binding protein, *CgCBP*) highly expressed in the black mantles. Three folds (inner fold, middle fold and outer fold) were found on the mantle edge of oyster, but only the outer fold has the same colour as the shell. Transcriptomic comparison indicated that the three folds of the mantle edge are functionally differentiated, and the *CgCBP* gene is ultra-highly expressed only in the outer fold. We obtained new black and white shell periostraca from the shell notching experiment and found their structural differences by scanning electron microscope. Chitin was successfully extracted and identified from the shell periostraca. Proteomic analysis revealed differences in protein composition between the two shell periostraca. Particularly, the black shell periostraca have more proteins related to melanin biosynthesis and chitin binding compared with the white ones, and melanin particles were observed in the black mantle edge using transmission electron microscope. Magnetic bead binding and Western blot experiments showed that the CgCBP protein can specifically bind to chitin and in vivo RNAi screening indicated that *CgCBP* knockdown can change the structure of the shell periostracum and reduce its pigmentation. All these results suggest that the outer fold of mantle may have more important roles in shell pigmentation, shell periostracum structure is functionally correlated with shell pigmentation, and the *CgCBP* gene ultra-highly expressed in the outer fold may influence shell pigmentation by affecting its periostracum structure.

## Introduction

Molluscs have colourful shells and shell patterns. Shell colour can be roughly categorised as pigment or structural and is mainly determined by the pigment produced by the mantle; the microstructure of the calcium carbonate crystal constituting the shell and the diet of the mollusc may also affect shell colour^1^. Mollusc shell pigments, including melanin^2,3^, porphyrins, protoporphyrins^4–6^ and carotenoids^7,8^, are mainly produced through biosynthesis or obtained through diet^6^. Pigments and pigment-related proteins in shells may affect colours and pigments distribution inside and outside shells^9^. Shell colour and pattern are phenotypic characteristics and important ecological traits of molluscs because of their role in communication, temperature regulation and survival behaviour in habitat^10, 11^. Hence, research on these characteristics contributes to evolutionary biology and population genetics^6,12^. Additionally, shell colour is heritable and is related to the growth, survival, immunity, antioxidant capacity and pearl colour of molluscs^13,14^. This feature can be selected as an economic trait to obtain new mollusc varieties^13,15,16^. Therefore, understanding the pigmentation mechanism of molluscs is important in investigating the evolutionary and ecological significance of shell colour and carrying out shell colour breeding.

As one of the most crucial economic varieties in aquaculture^17^, *Crassostrea gigas* is widely distributed and extensively studied due to its economic, biological and ecological importance^18^. Varieties of this species with various shell colours (white, gold and black) have been selected and bred to improve its commercial value^19,20^. Evidence from lncRNA, RNAi and other techniques proves that the shell mantle is an important tissue for shell colour^13,19,21,22^, however, the specific mechanism of oyster shell pigmentation remains unclear. Additionally, the outer surface of oyster has a radial stripe pattern except white, gold and black pure colours, and the striped pigmentation is a quantitative trait controlled by multiple genes^23^. In nature, striped patterns exist on the body surfaces of some fish and mammals, and the molecular mechanisms underlying their formation are different. As an interesting biological problem, stripe formation has been extensively and deeply studied^24–27^, but its mechanism in shells is rarely examined.

Many verification technologies for gene function cannot be applied in molluscs, and the heterozygosity of the oyster genome is extremely high^18,20^. Hence, screening out reliable trait-related genes by reasonable experimental design is especially important for oyster research. In the present study, *C. gigas* with black and white stripes on shell surface was selected to eliminate the interference caused by differences in individual genetic backgrounds and accurately screen out the genes related to shell colour. The black and white mantles of the same individual were paired and compared for first clue. Then transcriptomic and proteomic analyses, scanning electron microscopy, in vitro protein expression analysis, Western blot, chitin-binding assay, RNA interference and other methods were adopted to investigate the mantle and shell periostracum of black-, white- and striped-shelled oysters for understanding the pigmentation mechanism of shell.

## Results

### The colour of the shell stripes is corresponding to that of the mantle edge, and the *CgCBP* gene is highly expressed in the black mantle edge

According to the observation statistics of 100 striped-shell oysters, the distribution of black and white stripes on the shell is highly correlated with that on the mantle edge. Additionally, the black and white stripes with radioactive distribution on the shell have a one-to-one correlation with those on the mantle edge (Fig. 1a).

**Fig. 1.**
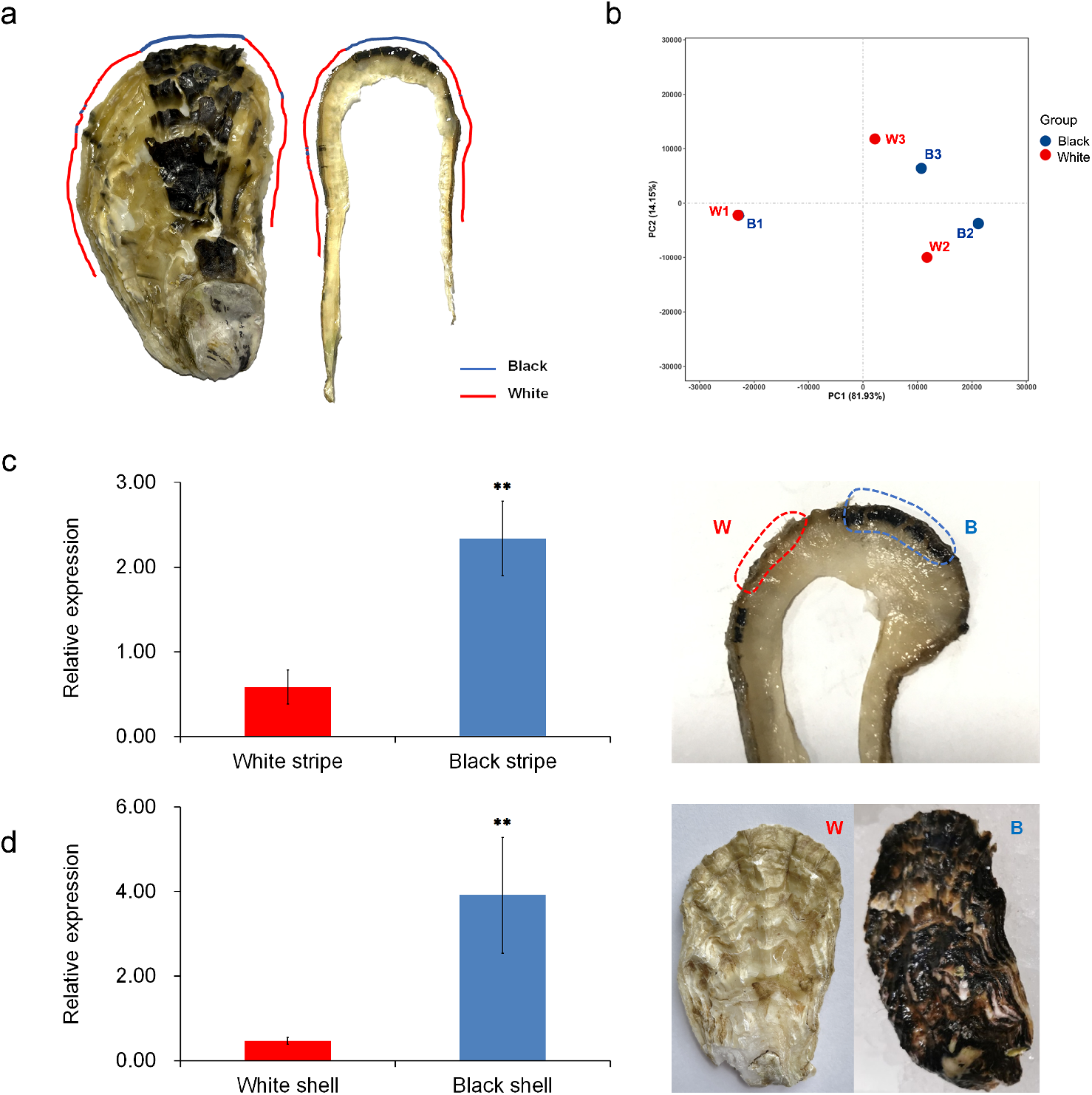
Physiological observation and transcriptome analysis of the mantle edge of *C. gigas.* (a) Shell stripes have a one-to-one correlation with mantle edge stripes (blue line: black stripe, red line: white stripe); (b) PCA analysis of striped mantle transcriptomes (blue point: black stripe, red point: white stripe); (c) Expression level of *CgCBP* gene in the black and white mantles of striped oysters (W: white stripe, B: black stripe); (d) Expression level of *CgCBP* gene in the mantles of black- and white-shelled oysters (W: white shell, B: black shell).

PCA analysis (Fig. 1b) on the transcriptomic data of the black and white striped mantles showed that the transcriptome of the black and white mantles of the same individual have a good clustering relationship. Nevertheless, significant differences in expression patterns are found among different individuals. Therefore, differential analysis was firstly conducted on the transcriptomic data of black and white mantles of the same specimen to exclude the interference of genetic background caused by individual differences. The presence of these differentially expressed genes between the black and white mantles of one or more oysters was then counted for predicting the reliability of these genes. A total of 90 differentially expressed genes are detected between the black and white mantle transcriptomes. However, most of these genes only appear in one individual, and only a few are found in the mantle of multiple individuals at the same time. Among these genes, LOC105336951 is significantly overexpressed in the black mantle of three striped individuals. Given that the function of LOC105336951 has not been annotated, its structural domain was analysed. The result revealed that this gene contains two chitin-binding domains (ChtBD2) and is henceforth named the *Crassostrea gigas* chitin-binding protein gene *(CgCBP).* RT-qPCR showed that the expression level of the *CgCBP* gene in the black mantle is significantly higher than that in the white mantle of striped-shell oysters (*p*<0.01) (Fig. 1c), and this finding further supports the difference analysis results of transcriptomic data. *CgCBP* gene expression was further verified in black- and white-shelled oysters, and RT-qPCR indicated that its expression level in the mantle is significantly higher in black oysters than in white ones (*p*<0.01) (Fig. 1d). According to the study on the mantle of striped shell, black- and white-shelled oysters, we further confirmed that the *CgCBP* gene is highly expressed in the black mantle.

### Among the three folds of mantle, only the outer fold has the same colour as the shell and ultra-high expression of the *CgCBP* gene

According to the previous observation for striped-shell oysters, the shell colour is highly correlated with its mantle edge colour. Observation statistics was carried out for 300 oysters with black and white shell colours to further clarify the relation between shell and mantle edge colours. The result showed that the correlation model of the shell colour and the colour of the mantle edge does not apply to all mantle edge structures. This relationship only exists in the outer fold, and the colours of the middle and inner folds have no correlation with the shell colour (Fig. 2a).

**Fig. 2.**
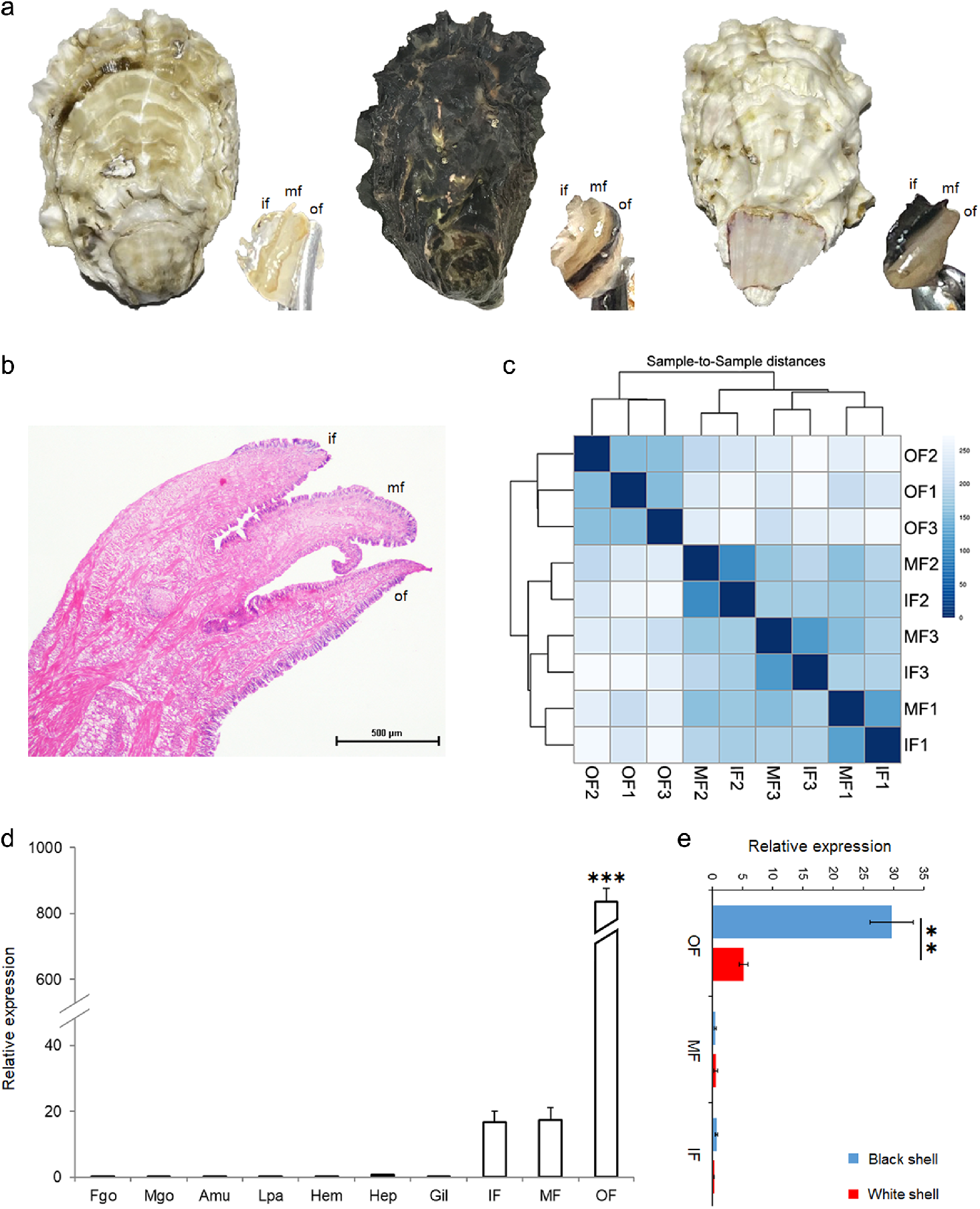
Physiological observation and transcriptome analysis of the three folds of mantle edge. (a) Relationship between the colour of the shell and that of three mantle folds: shell colour is only correlated with the colour of the outer fold but not with that of the inner and middle folds. if, inner fold; mf, middle fold; of, outer fold. (b) Paraffin sections of the three folds of the mantle edge were observed and stained with HE, bar=500 μm. (c) Clustering analysis of the transcriptome expression of the three folds. IF, inner fold; MF, middle fold; OF, outer fold. (d) *CgCBP* expression level in various tissues. Fgo, female gonad; Mgo, male gonad; Amu, adductor muscle; Lpa, labial palps; Hem, haemolymph; Hep, hepatopancreas; Gil, gill. (e) *CgCBP* expression level in the three folds of black- and white-shelled oysters.

The three folds of the mantle edge of three black-shelled oysters were extracted for transcriptomic analysis. According to the analysis on the nine transcriptomes, the outer folds of the three individuals have a good clustering relationship (Fig. 2c), and their expression patterns significantly differ from those of the inner and middle folds. The inner and middle folds have similar expression patterns and are clustered together. A total of 1034 genes (Supplementary Data 1) have significantly high expression in the outer fold. One of these genes is *CgCBP,* and its expression level in the outer fold is significantly higher than that in the middle and inner folds. RT-qPCR analysis was further conducted on the expression level of *CgCBP* in each fold of the mantle edge and other tissues. The result showed that the expression level of *CgCBP* in the outer fold of the black-shelled oysters is significantly higher than that in other tissues (*p*<0.001). This gene has a lower expression in the inner and middle folds than in the outer fold and is almost not expressed in the tissues of female and male glands, adductor muscle, labial palps, haemolymph, hepatopancreas and gill (Fig. 2d). Further RT-qPCR analysis on *CgCBP* expression in the three folds of the mantle of black- and white-shelled oysters revealed that this gene has significantly higher expression in outer fold of black-shelled oysters than in white-shelled oysters (Fig. 2e).

GO analysis (Supplementary Fig. 1a, Supplementary Data 2) showed that the differentially expressed genes with significantly high expression in the outer fold are mainly enriched with the chitin catabolic process, N-acetyl-beta-D-galactosaminidase activity and other biological processes related to chitin metabolism, copper ion binding, haem binding and other binding processes. Meanwhile, the highly expressed genes of the middle fold are mainly enriched with ossification involved in bone maturation, calcium ion binding, inner ear receptor cell stereocilium organisation, photoreceptor inner segment and other biological functions. The highly expressed genes of the inner fold are mainly enriched with a neuropeptide signalling pathway, calcium ion binding and other functions. KEGG analysis (Supplementary Fig. 1b, Supplementary Data 2) revealed that the highly expressed genes of the outer fold are mainly enriched with glycosphingolipid biosynthesis, glycosaminoglycan degradation, amino sugar and nucleotide sugar metabolism and other functions. The highly expressed genes of the inner and middle folds are mainly enriched with neuroactive ligand–receptor interaction.

### Black and white shell periostraca show differences in their electron microscope structures

After shell notching, the mantle became closely attached to the edge of the notch position. With the growth of the new shell periostraca, the mantle edge extended out from the notch position and tightly binds to the new shell periostraca (Supplementary Fig. 2a). At 0–36 h after cutting the shell, a thin and transparent periostracum began to grow at the notch position (Supplementary Fig. 2b). At 36– 72 h after cutting the shell, the thickness of the regenerated periostraca increased with its growth and forward extension. At the notch position of the striped shell, pigmentation could be found in the shell periostraca corresponding to the black outer fold, but not in those corresponding to the white outer fold (Supplementary Fig. 2c). At the notch positions of black- and white-shelled oysters, pigmentation could be observed in the periostraca of black-shelled oysters (Fig. 3a) but not in those of white-shelled oysters (Fig. 3d). At 72 h after shell notching, the regeneration periostraca continued to grow and gradually calcified.

**Fig. 3.**
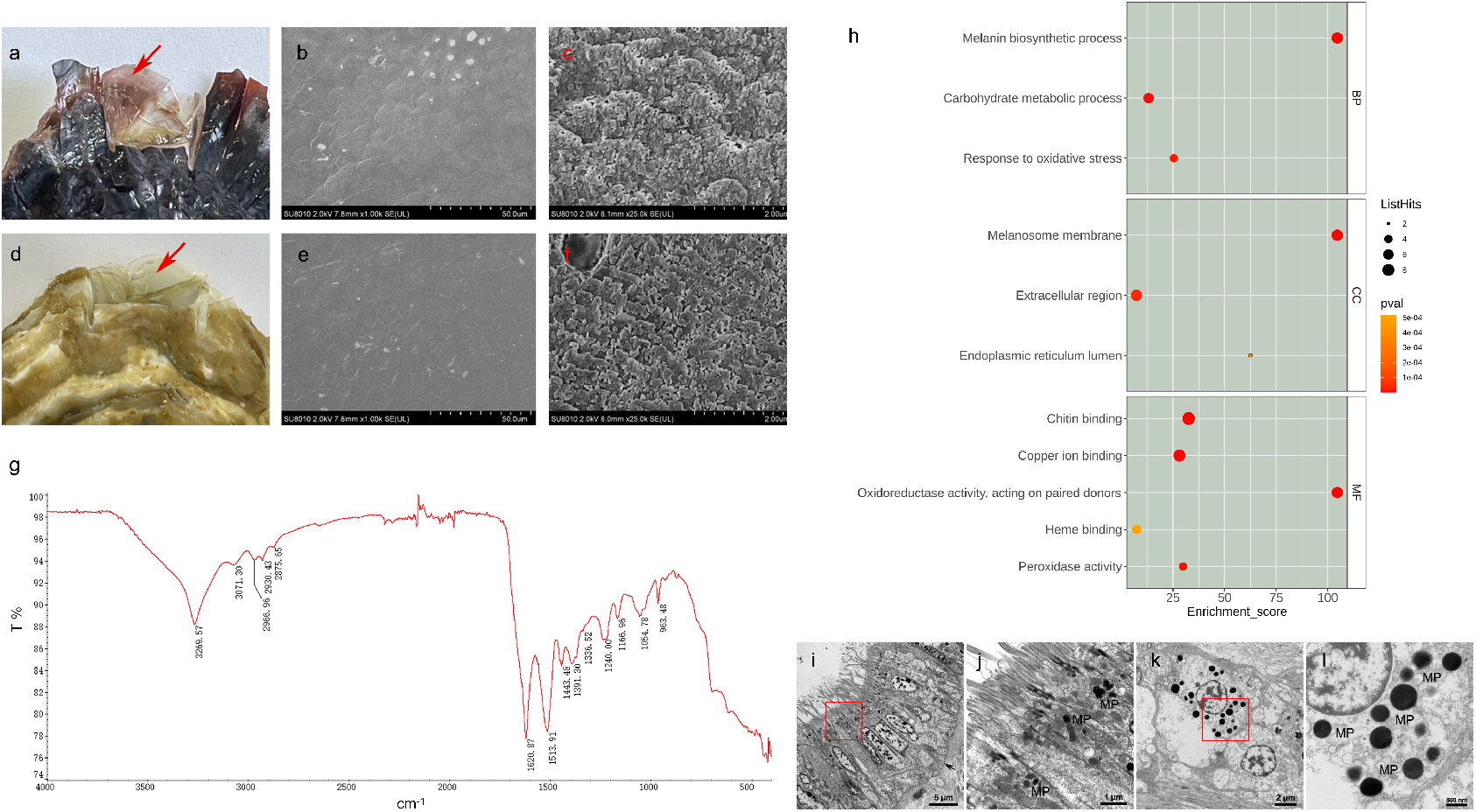
SEM of the regenerated periostraca after shell notching, chitin extraction, proteomic analysis and TEM of black mantle. (a, d) Regenerated periostraca of the black- and white-shelled oysters (the arrow shows regenerated periostracum, only the periostracum of the black-shelled oyster has pigmentation); (b, e) Outer surface structures of black and white periostraca are consistent; (c, f) Inner surface structures of black and white periostraca are different, lamellae of black periostracum (c) are larger than those of white periostracum (f); (g) Identification of chitin extracted from the regenerated periostraca by FTIR; (h) GO enrichment analysis of differentially expressed proteins in black and white periostraca; (i) Melanin particles (MP) in mantle epithelium, the arrow shows the basement membrane; (j) The enlarged area in the red box corresponding to Figure i; (k) MP in the cell located below the basement membrane; (l) The enlarged area in the red box corresponding to Figure k.

Phenotypically, pigmentation differs between black and white shell periostraca. Here a scanning electron microscope was used to determine differences in their microstructure. SEM image showed that the structures of the regenerated black and white shell periostraca on the outer surface were consistent and had no significant difference (Figs. 3b and 3e). However, significant differences were found in the electron microscopic structures on the inner surface (contact surface with the mantle) of the black and white shell periostraca. The lamella of the black shell periostraca is large, and the distance between two adjacent lamellae is approximately 1 μm. Many pore structures can be found in the lamellar edge (Fig. 3c). The lamellae of the white shell periostraca were small, and the distance between two adjacent lamellae was approximately 0.2–0.3 μm. The electron microscopic structure of the inner surface of the white shell periostraca was more fragmented (Fig. 3f).

### Chitin extraction and identification from the shell periostraca

Chitin powder was successfully extracted from the shell periostraca of *C. gigas.* FTIR identified N-H characteristic absorption peak at 3270 cm^-1^ and three characteristic absorption peaks of chitin (2962, 2930 and 2885 cm^-1^) in the range of 2800–3000 cm^-1^^28,29^(Fig. 3g). Therefore, chitin components are present in the oyster shell periostraca.

### The different protein composition between black and white shell periostraca and the observation of pigment particles in the mantle edge using TEM

A total of 309 high-quality proteins were identified by LC-MS/MS analysis. Among which, 262 and 219 proteins are found in black and white periostraca, respectively, and 172 proteins are present in both periostraca (Supplementary Data 3). Relative protein content analysis revealed that the highly expressed proteins in the black shell periostraca were enriched in 11 GO terms (Supplementary Table 4), including three functions related to biological process (melanin biosynthesis process, carbohydrate metabolism process, and response to oxidative stress), three functions related to cell components (melanosome periostraca and endoplasmic reticulum) and five functions related to molecular function (oxidoreductase activity, chitin binding, copper ion binding, peroxidase activity and haem binding) (Fig. 3h). Additionally, tyrosinase related to melanin biosynthesis (LOC105318244, LOC105320007, LOC105344040, LOC105328544, LOC105332955, LOC105324830), chitinase related to chitin binding and proteins containing chitin-binding domain (LOC105332577, LOC105332575, LOC105332622, LOC105345528, LOC105327512, LOC105336368, LOC105345527, LOC105332576) were highly expressed in the black shell periostraca. Using TEM, we also found many electron-dense components of different sizes in the epidermis (Fig. 3i, 3j) and in some cells below the basement membrane (Fig. 3k, 3l) of the black mantle edge. These electron-dense components were speculated to be melanin particles (MP). These cells located below the basement membrane with melanin particles were speculated to be melanocytes. Among these cells, melanosomes with different electron-density components and irregularly shaped membranes were distributed in the cytoplasm (Fig. 3l).

### CgCBP protein can specifically bind to chitin

According to the structural domain prediction of the amino acid sequence of CgCBP protein, a signal peptide was found at its N-terminal, followed by two chitin-binding domain 2 (ChtBD2) (Fig. 4a). Given that this signal peptide indicated that the CgCBP protein should be secrete protein, it was removed in the reconstructed plasmid that was expressed in vitro to verify whether ChtBD2 can bind with chitin. Western blot showed that the CgCBP protein and negative control protein are successfully expressed in HEK293T cells. After binding with chitin magnetic beads, some of the CgCBP protein and negative control protein that were not bound with chitin magnetic beads were eluted in the first washing solution. No signal of these proteins appeared in the fifth washing solution, indicating that the chitin magnetic beads were completely washed. In the final protein eluent, only the signal band for CgCBP protein appeared, indicating that it could specifically bind to chitin magnetic beads (Fig. 4b). This experiment showed that the CgCBP protein had a chitin binding ability.

**Fig. 4.**
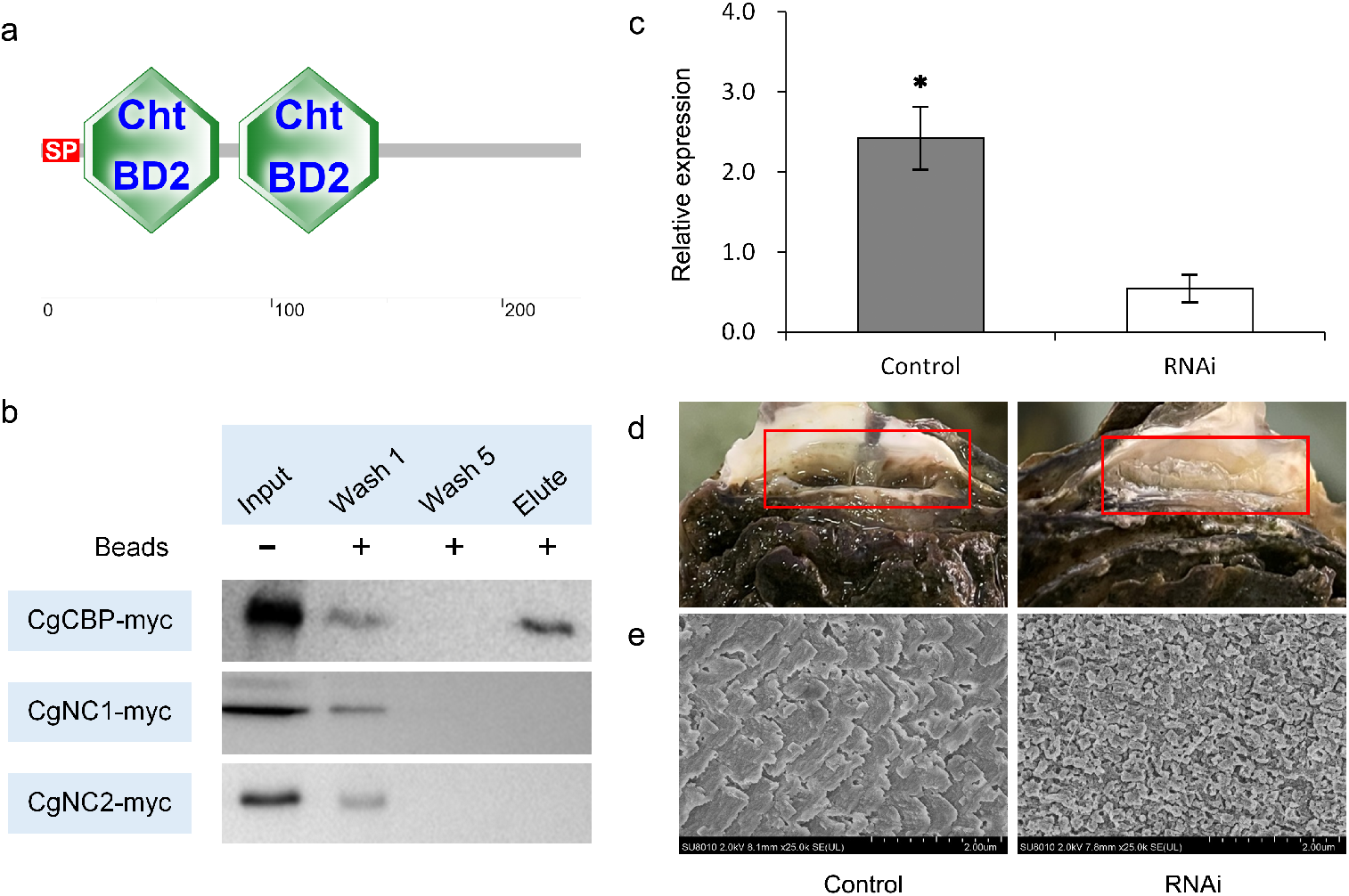
Verification of the chitin-binding function of CgCBP protein and RNAi of *CgCBP* in black-shelled oysters. (a) CgCBP protein was predicted to contain a signal peptide (SP) and two ChtBD2 domains; (b) Western blot results of the binding ability of CgCBP-myc recombinant protein to chitin magnetic beads (WB: anti-myc, CgNC1-myc and CgNC2-myc: negative control proteins); (c) *CgCBP* expression level was significantly down-regulated after RNAi; (d) The regenerated periostraca after RNAi have almost no pigmentation and grow slowly (the red boxes show the regenerated periostraca, the left is the control group and the right is the RNAi group); (e) SEM structure of the regenerated periostraca after RNAi (the right) is more irregular and incomplete than the control group (the left).

### *CgCBP* gene knockdown results in structural changes in the shell periostracum and reduces its pigmentation

CgCBP-dsRNA was injected into oysters for RNAi to investigate the physiological function of *CgCBP.* RT-qPCR results after RNAi showed that the relative expression level of *CgCBP* in the outer fold of the RNAi group was significantly decreased (p<0.05) (Fig. 4c). Therefore, CgCBP-dsRNA could significantly inhibit the mRNA expression level of *CgCBP* in the outer fold of the mantle. Observation of the growth of the shell periostraca at the notch position revealed that the regenerated shell periostraca in the RNAi group had almost no pigmentation, and the shell periostraca structure was irregular and grew slowly. By contrast, the periostraca in the control group had pigmentation, its edge and surface were smooth, and its growth was faster than that in the RNAi group (Fig. 4d). SEM was used to further compare the electron microscopic structures of the two periostraca. In the control group, the lamellar structure of periostraca was orderly arranged and smooth in edges, and its shape resembled a clear flat plate. The lamella was in a ladder shape with a relatively loose arrangement. In the RNAi group, the inner surface structure of the periostraca was destroyed and exhibited an irregular arrangement and incomplete shape (Fig. 4e).

## Discussion

### Innovation and advantage of experimental design

In the previously study of shell colour, oysters with black shell and white shell were usually chosen for the analysis of transcriptomic differences between black and white mantles^12,30^. In this study, the black and white mantles of the same striped individual were selected for pair analysis (Fig. 1) to maximumly eliminate the influence of differences in the individual genetic background and increase the reliability of the screened genes associated with shell colour. Furthermore, adjacent black and white mantles were selected to eliminate the interference caused by differences in tissue locations. Finally, the number of repeats of the differentially expressed genes between black and white mantles in three striped individuals were counted, and the genes with significant differences in multiple individuals at the same time were screened out to ensure the reliability of results. So, comparing different colour mantle edges of the same striped individual may be a better choice for shell colour research in molluscs.

### The three folds of mantle edge are functionally differentiated, and only the outer fold plays a key role in shell colour formation

Mantle plays an important role in shell biological mineralisation^31–33^ and is mainly divided into central mantle and mantle edge. The central mantle is involved in the calcification and thickening of the shell, such as inhibiting CaCO3 crystal and regulating the growth of epithelial cells and Ca^2+^; the mantle edge contains a large number of conchiolin proteins prompting the shell to grow outward and participating in shell periostraca formation^18,32,34,35^. Shell colour is also correlated with the colour of the mantle edge^13,21,36,37^, some pigmentation-related genes are expressed in the mantle edge^38–41^. The mantle edge has outer, middle and inner folds^42,43^. However, previous studies only focussed on the entire mantle edge, and the specific functions and differences of its three folds were not thoroughly examined. In the present work, detailed biological observations and statistics were conducted on oysters with black, white and striped shells. The findings showed that oyster shell colour is only correlated to the colour of the outer fold of the mantle edge.

Transcriptomic analysis was conducted to compare the functional differences among the three folds. The result shows that the functional differentiation between the inner fold and middle fold is relatively small; their functions are more reflected in calcium ion binding, neuropeptide signal regulation, sensitisation and other functions. However, many proteins encoding the chitin-binding domain in the highly expressed genes of the outer fold are enriched in the chitin catabolic process, N-acetyl-beta-D-galactosaminidase activity and other biological processes. Proteins with chitin-binding domains play an important role in the shell biological mineralisation and shell structural composition and can bind with calcium and chitin to participate in shell growth^44–49^. As an important component of the shell skeleton, chitin acts as a coordinating switch between the forming shell and the cells^50^ and plays an important role in shell biological mineralisation^51,52^. In addition, N-acetyl-beta-D-galactosaminidase activity can degrade chitin and remodel its scaffold^53^. Therefore, the function of the outer fold may be mainly reflected in the formation of the chitin framework and its participation in biological mineralisation. The corresponding correlation between the colour of the outer fold and shell (Fig. 2a) suggests a certain correlation between chitin and shell colour.

### Shell periostraca structure is functionally correlated with its pigmentation

Chitin and structural proteins are abundant in cuttlefish’s beak, butterfly’s wings and silkworm’s epidermis^54–57^. Chitin cross-links with structural proteins to form a network scaffold structure and provide a platform for melanin; however, pigmentation changes with the structure of the beak, wing and epidermis^54,55,57^. Melanin can be found in chitin cross-linked structures in perisarc, beak and other tissues of marine invertebrates^56,58^, and chitin–melanin complex can be extracted from insects^59^. As *C. gigas,* Moon scallop *Amusium pleuronectes* belongs to bivalves, and in the study of the differential genes between red and white shell moon scallops, some framework macromolecules genes, including several chitin-related genes, were found^38^. All these previous findings indicate that the chitin-rich structures of invertebrates may have a vital relationship with pigmentation.

According to the results of the shell notching experiment, pigments have been deposited before the regenerated periostraca has accumulated a large amount of calcium to become “shells”, and the pigmentation of shell periostraca only occurs in the black-shelled oysters and the black striped part of the striped oysters (Fig. 3a, Supplementary Fig. 2c). In addition to different pigmentation conditions, significant differences in the size and morphology of the lamellar structure were found between the electron microscopic structures of the black and white periostraca (Figs. 3c and 3f). This finding suggests that the shell periostraca structure could affect pigmentation. Proteomic analysis of black and white shell periostraca showed that the black periostraca contains some tyrosinases and proteins with chitin-binding domains that are enriched in the GO terms of melanin biosynthesis and chitin binding (Fig. 3h, Supplementary Table 4). The proteins related to melanin biosynthesis and chitin binding proteins simultaneously appear in the differential protein set of black and white periostraca, which further suggests that the pigmentation is functionally correlated with chitin in the shell periostraca.

### *CgCBP* gene may influence pigmentation by affecting the shell periostraca structure

According to the transcriptomic and RT-qPCR data, the *CgCBP* gene is significantly overexpressed in the black mantle edge whether in striped-shell oysters (Fig. 1c) or black-shelled oysters (Fig. 1d), the expression level of this gene is significantly higher in the outer fold of black-shelled oysters than white-shelled ones (Fig. 2e). Particularly, compared with other tissues, *CgCBP* gene is ultra-highly expressed in the outer fold (Fig. 2d). It is also found that only the outer fold has the same colour as the shell among the three folds of mantle (Fig. 2a). These results indicate that the *CgCBP* gene is associated with the shell pigmentation of *C. gigas.*

CgCBP has two ChtBD2 domains. The magnetic bead binding experiment confirmed that the CgCBP protein could specifically bind to chitin (Fig. 4). Additionally, the oyster shell periostracum contains a large amount of chitin (Fig. 3g). The mantle edge is the main organ that forms the shell periostraca^18,32,34^, and the *CgCBP* gene is only expressed in the outer fold of the mantle edge in *C. gigas* (Fig. 2d). Hence, this protein may be involved in the formation of the chitin framework. After knockdown of *CgCBP* by RNAi, the expression of this gene is significantly decreased, the structure of the shell periostraca is significantly changed, and melanin pigmentation is reduced (Fig. 5). In the notching experiment without RNAi, the white periostracum grew from the notch position of white shell (Fig. 3), its structure and pigmentation were similar to the RNAi group. Therefore, the *CgCBP* gene may influence pigmentation by affecting the shell periostracum structure in *C. gigas.*

## Methods

### Physiological observation and transcriptomic analysis of striped-shell oyster

A total of 100 oysters with black and white stripes on the shell surface were purchased from Yantai, Shandong Province, China and randomly selected for dissection. Stripes distribution on the shell surface and the corresponding mantle edge of each sample was observed and counted.

Three oysters with striped shells were selected, and the black and white mantle edges corresponding to the colour of the shell stripes were chosen for RNA-seq sequencing. For differential expression analysis, genes with differential expression of black and white mantles in the same individual were firstly screened out, and their occurrence in one or more individuals was counted. Raw data (raw reads) were processed using Trimmomatic^60^ and filtered to remove the reads containing poly-N or low-quality reads. Then, the resulting high-quality (HQ) reads were mapped to the oyster genome using hisat2^61^. The read counts and FPKM^62^ were calculated using htseq-count^63^ and cufflinks^64^. DEGs were identified using DESeq package^65^ with *P value* < 0.05 and foldchange >2 as threshold. GO enrichment and KEGG^66^ pathway enrichment analysis of DEGs were performed using R.

Twelve striped-shell oysters were randomly selected, and their black and white mantle edges were collected and frozen in liquid nitrogen and stored at −80°C to determine the expression of the differential genes screened from the transcriptomes. The total RNA was extracted using RNAiso Plus Reagent (Takara, Japan). The integrity of RNA was checked by agarose gel electrophoresis. The first-strand cDNA was synthesized using an AGEvo M-MLV Plus cDNA Synthesis Kit (Accurate Biology, China) using treated RNA (1.0 μg) as a template and oligo(dT)-adaptor as a primer. The cDNA synthesis reaction was performed at 42 °C for 30 min and terminated at 95 °C for 5 min. Primers for qPCR are given in Supplementary Table 1. Real-time qPCR was conducted on the Prime Pro 48 real-time qPCR system (Techne, UK). The 2^-ΔΔt^ method was used to calculate the relative expression of target genes. Gene expression differences between black and white stripe mantle edge were compared using paired-samples t tests. Differences were considered significant if *p* < 0.05.

Eight black-shelled and eight white-shelled oysters were randomly selected, and their mantle tissues were collected to further verify whether the expression pattern differences detected in striped-shell oysters can also be observed in black- and white-shelled oysters. The methods for RNA extraction and RT-qPCR are described as above. Gene expression differences between black- and white-shelled oyster were compared using Student’s t tests. All data were analyzed using SPSS 21.0 (IBM Corp, Armonk, USA).

### Physiological observation and transcriptomic analysis of the three folds of the mantle edge

A total of 300 oysters with black and white shell colours purchased from Yantai, Shandong Province, China, were randomly selected for dissection. The colours of the shell and the three folds (outer, middle and inner folds) of the mantle edge of each oyster were observed and counted.

Three black-shelled oysters were randomly selected to further study the relationship between the three folds of the mantle edge and the black shell colour traits. RNA-seq transcriptomic sequencing was performed on the outer, middle and inner folds of the mantle edge, and gene expression differences among different folds were analysed. The sequencing and analysis methods for the library are same as described above. qPCR was used to verify the gene expression pattern in the three folds and other tissues. Additionally, three black-shelled oysters were randomly selected, and their female and male gonads, adductor muscle, labial palps, haemolymph, hepatopancreas, gills, outer, middle and inner folds of the mantle edge were extracted, frozen in liquid nitrogen and stored at −80°C for future use. Seven black- and seven white-shelled oysters were randomly selected for dissection to determine the expression of genes in the three folds of black- and white-shelled oysters. The outer, middle and inner folds of the mantle edge were obtained for quantitative analysis. The methods for RNA extraction and RT-qPCR are same as described above.

### Black and white shell notching and scanning electron microscopy (SEM) observation

One-year-old black and white-shelled Pacific oysters *C. gigas* were obtained from Yantai (Shandong Province, China) in January 2021. After collection, the oysters were acclimated at 15°C in aerated seawater for 3 days, then 10 black-shelled and 10 white-shelled individuals were randomly chosen for notching. A fan-shaped notch with length and width of 15 × 5 mm was cut in the posterior of the right shell edge. The regeneration shell periostracum which begun at the notching site were dissected, immediately frozen in liquid nitrogen, and stored at −80°C.

The regenerated periostraca of black- and white-shelled oysters were dissected, rinsed with deionized water immediately and then air-dried at room temperature. The well-dried shell periostraca were cut into pieces of 1-2 mm by mechanical means, and then coated with gold by using the ion sputtering system. SEM (HITACHI SU8010) was used to observe the microstructures of the samples.

### Chitin extraction and Fourier transform infrared spectroscopy (FTIR)

The regenerated periostraca after shell notching were separated, cleaned and dried, soaked in 2% hydrochloric acid solution for 30–60 min for demineralisation, treated with 2 mol/L NaOH solution for 6 h to remove proteins, bleached and decolourised with 10% (v/v) H_2_O_2_ solution, rinsed with deionised water and dried at 60°C to obtain the chitin powder. A small amount of chitin powder was used to cover the diamond sheet test table, and the infra-red spectrum within 500–4000 cm^-1^ was determined at room temperature.

### Protein preparation and LC-MS/MS analysis and transmission electron microscopy (TEM) observation of the mantle edge

Six black-shelled and white-shelled oysters were obtained and notched. The regenerated periostraca of black and white-shelled oysters were dissected. The total proteins were isolated from regenerated shell periostracum and then digested with sequencing grade Trypsin (HLS TRY001C) incubated at 37°C for 16 h. The tryptic peptide digests were desalted and vacuum dried. The samples were then analyzed using UltiMate 3000 RSLCnano HPLC system connected to mass spectrometer. The sample was loaded onto columns at a flow rate of 300 nl/min. The sequential separation of peptides on trap column (100 μm × 20 mm, RP-C18, Agilent) and analysis column (75 μm × 150 mm, RP-C18, New Objective) was accomplished using a segmented gradient from Solvent A (0.1% formic acid in water) to 95% Solvent B (0.1% formic acid/50% ACN in water) for 95 min. The column was re-equilibrated to its initial highly aqueous solvent composition before each analysis.

The mass spectrometer was operated by Q-Exactive plus (Thermo Fisher Scientific) and acquired over a range of 350-2000 m/z. The resolving powers of the MS scan and MS/MS scan at 200 m/z for the Orbitrap Elite were set as 70,000 and 17,500, respectively. The top 20 most intense signals in the acquired MS spectra were selected for further MS/MS analysis. The ions were fragmented through higher energy collisional dissociation with normalized collision energies of 28 eV. The maximum ion injection times were set at 100 ms for the survey scan and 50 ms for the MS/MS scans, and the automatic gain control target values were set to 3.0e6 for full scan modes and 1e5 for MS/MS. The dynamic exclusion duration was 25s. The data were analyzed by software ProteomeDiscover 2.1, using the uniprot-(taipingyangmuli) Crassostrea gigas [29159] database. The MS1 tolerance was 10 ppm and MS2 tolerance was 0.05 Da, with the missed cleavage were set as 2. The static modification was Carbamidomethyl (C), and the dynamic modification was set as Acetyl (Protein N-term), Deamidated (NQ), Oxidation (M).

The mantle edge samples were obtained from black-shelled oysters, fixed with glutaraldehyde for 4 h and postfixed with 1% osmium tetroxide for 1 h. The tissue blocks were dehydrated with 50%, 70%, 90%, and 100% acetone for 15 min each, embedded in Epon 812 resin and cured at 37°C, 45 °C, and 60°C for 24 h each. After determining the research areas by semi-thin sections and toluidine blue staining, the samples were cut into ultrathin sections of 70 nm with ultratome (Reichert-Jung ULTRACUT E, Austria). The samples were stained with uranium acetate and lead citrate double staining and observed using TEM (JEOL, JEM1200, Japan).

### Validation of the chitin-binding function of CgCBP protein

To construct protein-expressing plasmids, Phusion High-Fidelity DNA Polymerase (Thermo Scientific, USA) was used to amplify the gene fragments with the designed primers (given in Supplementary Table 2). Primers CgCBP-myc-F and CgCBP-myc-R were used for amplification of the ORF (signal peptide removed) of CgCBP. Another 3 proteins with no chitin-binding domain were chosen as negative control (Supplementary Table 2). The pCMV-myc plasmid was digested with EcoRI (New England Biolabs). The PCR products and digested plasmids were electrophoresed, isolated, and purified using the gel purification kit (Sangon, China). After purification, the PCR products and digested plasmids were fused into complete expression plasmids using the Ligation-Free Cloning System (Applied Biological Materials, Inc., Canada).

Due to the lack of marine mollusc cell lines, HEK293T cells were used for protein expression. Dulbecco’s modified Eagle’s medium–high glucose (DMEM) medium (HyClone, USA) was used for cell culture. The DMEM medium was supplemented with 5% penicillin/streptomycin resistance solution (Sangon, China) and 10% fetal bovine serum (Gibco, USA). The cells were grown at 37 °C with 5% CO_2_ and sub-cultured every 2–3 days. Before transfection, HEK293T cells were cultured into two 10 cm Petri dishes (Corning, USA) and cultured for 24 h.

The myc-tagged plasmids (CgCBP-myc and negative control CgNC-myc) were then transfected into the cells. After 24 h, the HEK293T cells were collected in a 1.5 ml tub and lysed with 700 ul cell lysis buffer (Beyotime, China) and 7 ul protease inhibitor (Beyotime, China) for 30 min on ice. 50 ul chitin magnetic beads (Sigma, USA) were well-washed by column binding buffer (500 mM NaCl, 20 mM Tris-HCl, 1 mM EDTA, 0.05% Tween-20, pH 8.0 @ 25°C) and added to the cell lysates. The mixture was incubated at 4°C and gently shaken for 2 h. After incubating, the beads were well-washed by column binding buffer 5 times and then incubated in 2× Protein SDS PAGE Loading (Takara) 100°C for 5 min to completely elute the proteins. The 5 times washing supernatant and the final eluting supernatant were separately collected for western blot.

### In vivo verification of *CgCBP* function by RNA interference (RNAi)

Six black-shelled oysters were subjected to double-stranded RNA (dsRNA) mediated silencing of *CgCBP* gene and 6 ones were as control. Before the experiments, all oysters were shell notched following the method described in “black and white shell notching” experiment. Before and during the experiments, all oysters were acclimated in aerated tanks at 15°C and fed with golden algae. CgCBP dsRNA and NC-dsRNA (species-specific sequence in *Caenorhabditis elegans*) were synthesized by Sangon, China. Sequences of dsRNA are given in Supplementary Table 3. The synthesized dsRNA was diluted to 0.8 μg/μl with nuclease-free 1×PBS. For each oyster, 100 μl CgCBP dsRNA or 100 μl NC dsRNA was injected into the muscle for 8 days. Then, the mantle edge was dissected for *CgCBP* expression analysis by RT-qPCR and the shell periostracum was dissected for color observation and SEM.

## Data availability

The raw sequencing data for *C. gigas* mantle RNA-seq have been deposited in NCBI Sequence Read Archive under the accession numbers of SRR17994559, SRR17994565, SRR17994566, SRR17994567, SRR17994568, SRR17994569, SRR17994570, SRR17994571, SRR17994572, SRR17994573, SRR17994574, SRR18490888, SRR18490892, SRR18490894, SRR18490896. The proteomic data have been deposited in PRIDE database under the accession number PXD033919. The authors declare that the main data supporting the findings of this study are available within the article and its Supplementary Information, further inquiries can be directed to the corresponding authors.

## Acknowledgements

We acknowledge grant support from the National Natural Science Foundation of China (Nos. 41876193, 42076088, and 41906088), National Key R&D Program of China (No. 2018YFD0901400), the Fine Agricultural Breeds Project of Shandong Province, China (2019LZGC020), the Shandong Provincial Natural Science Foundation, China (No.ZR2019MC002), the Special Funds for Taishan Scholars Project of Shandong Province, China (No. tsqn201812094), the Modern Agricultural Industry Technology System of Shandong Province, China (SDAIT-14-03), and the Plan of Excellent Youth Innovation Team of Colleges and Universities in Shandong Province, China (2019KJF004).

## Author contributions

X.W. and M.Z. conceived and designed the study. M.L. and J.T. conducted experiments and data analysis. M.Z. and M.L. conducted RNA-seq experiments and data analysis. B.H., Y.L. and X.S. contributed to western blot experiments. L.W. and Y.H. contributed to TEM analysis. M.Y. and XN.W. contributed to shell notching experiments and SEM analysis. N.F., S.C. and Y.Z. contributed to oyster materials and experiments. X.W., M.Z. and M.L. wrote the paper. All authors approved the final manuscript.

## References

1. Williams S (2017). Molluscan shell colour. Biol. Rev. Camb. Philos. Soc. 92, 1039–1058. 10.1111/brv.12268.

2. Hao S, Hou X, Wei L, Li J, Li Z, and Wang X (2015). Extraction and identification of the pigment in the adductor muscle scar of pacific oyster *Crassostrea gigas*. PLoS One 10, e0142439. 10.1371/journal.pone.0142439.

3. Sun X, Liu Z, Zhou L, Wu B, Dong Y, and Yang A (2016). Integration of Next Generation Sequencing and EPR Analysis to Uncover Molecular Mechanism Underlying Shell Color Variation in Scallops. PLoS One 11, e0161876. 10.1371/journal.pone.0161876.

4. Verdes A, Cho W, Hossain M, Brennan P L, Hanley D, Grim T, Hauber M E, and Holford M (2015). Nature’s Palette: Characterization of Shared Pigments in Colorful Avian and Mollusk Shells. PLoS One 10, e0143545. 10.1371/journal.pone.0143545.

5. Bonnard M, Cantel S, Boury B, and Parrot I (2020). Chemical evidence of rare porphyrins in purple shells of *Crassostrea gigas* oyster. Sci. Rep. 10, 12150. 10.1038/s41598-020-69133-5.

6. Saenko S, and Schilthuizen M (2020). Evo-devo of shell colour in gastropods and bivalves. Curr. Opin. Genet. Dev. 69, 1–5. 10.1016/j.gde.2020.11.009.

7. Liu H, Zheng H, Zhang H, Deng L, Liu W, Wang S, Meng F, Wang Y, Guo Z, Li S, et al. (2015). A de novo transcriptome of the noble scallop, *Chlamys nobilis*, focusing on mining transcripts for carotenoid-based coloration. BMC Genomics 16, 44. 10.1186/s12864-015-1241-x.

8. Zhao L, Li Y, Li Y, Yu J, Liao H, Wang S, Lv J, Liang J, Huang X, and Bao Z (2017). A Genome-Wide Association Study Identifies the Genomic Region Associated with Shell Color in Yesso Scallop, *Patinopecten yessoensis*. Mar. Biotechnol. (NY). 19, 301–309. 10.1007/s10126-017-9751-y.

9. Grant H E, and Williams S T (2018). Phylogenetic distribution of shell colour in Bivalvia (Mollusca). Biol. J. Linn. Soc. 125, 377. 10.1093/biolinnean/bly122.

10. Lindberg D R, and Pearse J S (1990). Experimental manipulation of shell color and morphology of the limpets *Lottia asmi* (Middendorff) and *Lottia digitalis* (Rathke) (Mollusca: Patellogastropoda). J. Exp. Mar. Biol. Ecol. 140, 173–185. 10.1016/0022-0981(90)90125-V.

11. Sokolova I M, and Berger V J (2000). Physiological variation related to shell colour polymorphism in White Sea *Littorina saxatilis*. J. Exp. Mar. Bio. Ecol. 245, 1–23. 10.1016/S0022-0981(99)00132-X.

12. Feng D, Li Q, Yu H, Zhao X, and Kong L (2015). Comparative Transcriptome Analysis of the Pacific Oyster *Crassostrea gigas* Characterized by Shell Colors: Identification of Genetic Bases Potentially Involved in Pigmentation. PLoS One 10, e0145257. 10.1371/journal.pone.0145257.

13. Brake J, Evans F, and Langdon C (2004). Evidence for genetic control of pigmentation of shell and mantle edge in selected families of Pacific oysters, *Crassostrea gigas*. Aquaculture 229, 89–98. 10.1016/S0044-8486(03)00325-9.

14. Adzigbli L, Wang Z, Li J, and Deng Y (2020). Survival, retention rate and immunity of the black shell colored stocks of pearl oyster *Pinctada fucata martensii* after grafting operation. Fish Shellfish Immunol. 98, 691–698. 10.1016/j.fsi.2019.11.003.

15. Zou K, Zhang D, Guo H, Zhu C, Li M, and Jiang S (2014). A preliminary study for identification of candidate AFLP markers under artificial selection for shell color in pearl oyster *Pinctada fucata*. Gene 542, 8–15. 10.1016/j.gene.2014.03.029.

16. Wang C, Liu B, Liu X, Ma B, Zhao Y, Zhao X, Liu F, and Liu G (2017). Selection of a new scallop strain, the Bohai Red, from the hybrid between the bay scallop and the Peruvian scallop. Aquaculture 479, 250–255. 10.1016/j.aquaculture.2017.05.045.

17. Zhang G, Li L, Meng J, Qi H, Qu T, Xu F, and Zhang L (2016). Molecular Basis for Adaptation of Oysters to Stressful Marine Intertidal Environments. Annu. Rev. Anim. Biosci. 4, 357–381. 10.1146/annurev-animal-022114-110903.

18. Zhang G, Fang X, Guo X, Li L, Luo R, Xu F, Yang P, Zhang L, Wang X, Qi H, et al. (2012). The oyster genome reveals stress adaptation and complexity of shell formation. Nature 490, 49–54. 10.1038/nature11413.

19. Feng D, Li Q, Yu H, Kong L, and Du S (2018). Transcriptional profiling of long non-coding RNAs in mantle of *Crassostrea gigas* and their association with shell pigmentation. Sci. Rep. 8, 1436. 10.1038/s41598-018-19950-6.

20. Wang X, Xu W, Wei L, Zhu C, He C, Song H, Cai Z, Yu W, Jiang Q, Li L, et al. (2019). Nanopore Sequencing and De Novo Assembly of a Black-Shelled Pacific Oyster *(Crassostrea gigas)* Genome. Front. Genet. 10, 1211. 10.3389/fgene.2019.01211.

21. Xing D, Li Q, Kong L, and Yu H (2018). Heritability estimate for mantle edge pigmentation and correlation with shell pigmentation in the white-shell strain of Pacific oyster, *Crassostrea gigas*. Aquaculture 482, 73–77. 10.1016/j.aquaculture.2017.09.026.

22. Feng D, Li Q, and Yu H (2019). RNA Interference by Ingested dsRNA-Expressing Bacteria to Study Shell Biosynthesis and Pigmentation in *Crassostrea gigas*. Mar. Biotechnol. (NY). 21, 526–536. 10.1007/s10126-019-09900-2.

23. Xu C, Li Q, Yu H, Liu S, Kong L, and Chong J (2019). Inheritance of shell pigmentation in Pacific oyster *Crassostrea gigas*. Aquaculture 512, 734249. 10.1016/j.aquaculture.2019.734249.

24. Mallarino R, Henegar C, Mirasierra M, Manceau M, Schradin C, Vallejo M, Beronja S, Barsh G S, and Hoekstra H E (2016). Developmental mechanisms of stripe patterns in rodents. Nature 539, 518–523. 10.1038/nature20109.

25. Eom D S, and Parichy D M (2017). A macrophage relay for long-distance signaling during postembryonic tissue remodeling. Science 355, 1317–1320. 10.1126/science.aal2745.

26. Mallarino R, Pillay N, Hoekstra H E, and Schradin C (2018). African striped mice. Curr. Biol. 28, R299–R301. 10.1016/j.cub.2018.02.009.

27. Irion U, and Nusslein-Volhard C (2019). The identification of genes involved in the evolution of color patterns in fish. Curr. Opin. Genet. Dev. 57, 31–38. 10.1016/j.gde.2019.07.002.

28. Sudatta B P, Sugumar V, Varma R, and Nigariga P (2020). Extraction, characterization and antimicrobial activity of chitosan from pen shell, *Pinna bicolor*. Int. J. Biol. Macromol. 163, 423–430. 10.1016/j.ijbiomac.2020.06.291.

29. Cuong H N, Minh N C, Van Hoa N, and Trung T S (2016). Preparation and characterization of high purity beta-chitin from squid pens *(Loligo chenisis)*. Int. J. Biol. Macromol. 93, 442–447. 10.1016/j.ijbiomac.2016.08.085.

30. Wang J, Li Q, Zhong X, Song J, Kong L, and Yu H (2018). An integrated genetic map based on EST-SNPs and QTL analysis of shell color traits in Pacific oyster *Crassostrea gigas*. Aquaculture 492, 226–236. 10.1016/j.aquaculture.2018.04.018.

31. Sun X, Yang A, Wu B, Zhou L, and Liu Z (2015). Characterization of the mantle transcriptome of yesso scallop *(Patinopecten yessoensis):* identification of genes potentially involved in biomineralization and pigmentation. PLoS One 10, e0122967. 10.1371/journal.pone.0122967.

32. Bjarnmark N A, Yarra T, Churcher A M, Felix R C, Clark M S, and Power D M (2016). Transcriptomics provides insight into *Mytilus galloprovincialis* (Mollusca: Bivalvia) mantle function and its role in biomineralisation. Mar. Genomics 27, 37–45. 10.1016/j.margen.2016.03.004.

33. Johnson A B, Fogel N S, and Lambert J D (2019). Growth and morphogenesis of the gastropod shell. Proc. Natl. Acad. Sci. U. S. A. 116, 6878–6883. 10.1073/pnas.1816089116.

34. Yarra T, Blaxter M, and Clark M S (2021). A Bivalve Biomineralization Toolbox. Mol. Biol. Evol. 38, 4043–4055. 10.1093/molbev/msab153.

35. Yarra T, Ramesh K, Blaxter M, Huning A, Melzner F, and Clark M S (2021). Transcriptomic analysis of shell repair and biomineralization in the blue mussel, *Mytilus edulis*. BMC Genomics 22, 437. 10.1186/s12864-021-07751-7.

36. Kang J H, Kang H S, Lee J M, An C M, Kim S Y, Lee Y M, and Kim J J (2013). Characterizations of Shell and Mantle Edge Pigmentation of a Pacific Oyster, *Crassostrea gigas*, in Korean Peninsula. Asian-Aust. J. Anim. Sci. 26, 1659–1664. 10.5713/ajas.2013.13562.

37. Budd A, McDougall C, Green K, and Degnan B M (2014). Control of shell pigmentation by secretory tubules in the abalone mantle. Front. Zool. 11, 62. 10.1186/s12983-014-0062-0.

38. Huang R L, Zheng Z, Wang Q H, Zhao X X, Deng Y W, Jiao Y, and Du X D (2015). Mantle Branch Specific RNA Sequences of Moon Scallop *Amusium pleuronectes* to Identify Shell Color-Associated Genes. PLoS One 10, e0141390. 10.1371/journal.pone.0141390.

39. Ding J, Wen Q, Huo Z, Nie H, Qin Y, and Yan X (2021). Identification of shell-color-related microRNAs in the Manila clam *Ruditapes philippinarum* using high-throughput sequencing of small RNA transcriptomes. Sci. Rep. 11, 8044. 10.1038/s41598-021-86727-9.

40. Huang S, Jiang H, Zhang L, Gu Q, Wang W, Wen Y, Luo F, Jin W, and Cao X (2021). Integrated proteomic and transcriptomic analysis reveals that polymorphic shell colors vary with melanin synthesis in *Bellamya purificata* snail. J. Proteomics 230, 103950. 10.1016/j.jprot.2020.103950.

41. Zhu X, Zhang J, Hou X, Liu P, Lv J, Xing Q, Huang X, Hu J, and Bao Z (2021). A Genome-Wide Association Study Identifies Candidate Genes Associated With Shell Color in Bay Scallop *Argopecten irradians irradians*. Front. Mar. Sci. 8, 742330. 10.3389/fmars.2021.742330.

42. Zhang C, Xie L, Huang J, Chen L, and Zhang R (2006). A novel putative tyrosinase involved in periostracum formation from the pearl oyster (*Pinctada fucata*). Biochem. Biophys. Res. Commun. 342, 632–639. 10.1016/j.bbrc.2006.01.182.

43. Glover E A, and Taylor J D (2010). Needles and pins: acicular crystalline periostracal calcification in venerid bivalves (Bivalvia: Veneridae). J. Mollus. Stud. 76, 157–179. 10.1093/mollus/eyp054.

44. Kintsu H, Okumura T, Negishi L, Ifuku S, Kogure T, Sakuda S, and Suzuki M (2017). Crystal defects induced by chitin and chitinolytic enzymes in the prismatic layer of *Pinctada fucata*. Biochem. Biophys. Res. Commun. 489, 89–95. 10.1016/j.bbrc.2017.05.088.

45. Carini A, Koudelka T, Tholey A, Appel E, Gorb S N, Melzner F, and Ramesh K (2019). Proteomic investigation of the blue mussel larval shell organic matrix. J. Struct. Biol. 208, 107385. 10.1016/j.jsb.2019.09.002.

46. Jin C, Zhao J, Pu J, Liu X, and Li J (2019). Hichin, a chitin binding protein is essential for the self assembly of organic frameworks and calcium carbonate during shell formation. Int. J. Biol. Macromol. 135, 745–751. 10.1016/j.ijbiomac.2019.05.205.

47. Liao Z, Jiang Y T, Sun Q, Fan M H, Wang J X, and Liang H Y (2019). Microstructure and in-depth proteomic analysis of *Perna viridis* shell. PLoS One 14, e0219699. 10.1371/journal.pone.0219699.

48. Arroyo-Loranca R G, Hernandez-Saavedra N Y, Hernandez-Adame L, and Rivera-Perez C (2020). Ps19, a novel chitin binding protein from *Pteria sterna* capable to mineralize aragonite plates in vitro. PLoS One 15, e0230431. 10.1371/journal.pone.0230431.

49. Fan S, Zheng Z, Hao R, Du X, Jiao Y, and Huang R (2020). PmCBP, a novel poly (chitin-binding domain) gene, participates in nacreous layer formation of *Pinctada fucata martensii*. Comp. Biochem. Physiol. B Biochem. Mol. Biol. 240, 110374. 10.1016/j.cbpb.2019.110374.

50. Weiss I M, and Schonitzer V (2006). The distribution of chitin in larval shells of the bivalve mollusk *Mytilus galloprovincialis*. J. Struct. Biol. 153, 264–277. 10.1016/j.jsb.2005.11.006.

51. Falini G, Albeck S, Weiner S, and Addadi L (1996). Control of Aragonite or Calcite Polymorphism by Mollusk Shell Macromolecules. Science 271, 67–69. 10.1126/science.271.5245.67.

52. Falini G, Fermani S, and Ripamonti A (2002). Crystallization of calcium carbonate salts into beta chitin scaffold. J. Inorg. Biochem. 91, 475–480. 10.1016/s0162-0134(02)00471-3.

53. Merzendorfer H, and Zimoch L (2003). Chitin metabolism in insects: structure, function and regulation of chitin synthases and chitinases. J. Exp. Biol. 206, 4393–4412. 10.1242/jeb.00709.

54. Miserez A, Li Y, Waite J H, and Zok F (2007). Jumbo squid beaks: inspiration for design of robust organic composites. Acta. Biomater. 3, 139–149. 10.1016/j.actbio.2006.09.004.

55. Futahashi R, Shirataki H, Narita T, Mita K, and Fujiwara H (2012). Comprehensive microarray-based analysis for stage-specific larval camouflage pattern-associated genes in the swallowtail butterfly, *Papilio xuthus*. BMC Biol. 10, 46. 10.1186/1741-7007-10-46.

56. Tan Y, Hoon S, Guerette P A, Wei W, Ghadban A, Hao C, Miserez A, and Waite J H (2015). Infiltration of chitin by protein coacervates defines the squid beak mechanical gradient. Nat. Chem. Biol. 11, 488–495. 10.1038/nchembio.1833.

57. Qiao L, Yan Z W, Xiong G, Hao Y J, Wang R X, Hu H, Song J B, Tong X L, Che L R, He S Z, et al. (2020). Excess melanin precursors rescue defective cuticular traits in stony mutant silkworms probably by upregulating four genes encoding RR1-type larval cuticular proteins. Insect Biochem Mol Biol. 119, 103315. 10.1016/j.ibmb.2020.103315.

58. Hwang D S, Masic A, Prajatelistia E, Iordachescu M, and Waite J H (2013). Marine hydroid perisarc: a chitin- and melanin-reinforced composite with DOPA-iron(III) complexes. Acta. Biomater. 9, 8110–8117. 10.1016/j.actbio.2013.06.015.

59. Khayrova A, Lopatin S, and Varlamov V (2021). Obtaining chitin, chitosan and their melanin complexes from insects. Int. J. Biol. Macromol. 167, 1319–1328. 10.1016/j.ijbiomac.2020.11.086.

60. Bolger A M, Lohse M, and Usadel B (2014). Trimmomatic: a flexible trimmer for Illumina sequence data. Bioinformatics 30, 2114–2120. 10.1093/bioinformatics/btu170.

61. Kim D, Langmead B, and Salzberg S L (2015). HISAT: a fast spliced aligner with low memory requirements. Nat. Methods 12, 357–360. 10.1038/nmeth.3317.

62. Roberts A, Trapnell C, Donaghey J, Rinn J L, and Pachter L (2011). Improving RNA-Seq expression estimates by correcting for fragment bias. Genome Biol. 12, R22. 10.1186/gb-2011-12-3-r22.

63. Anders S, Pyl P T, and Huber W (2015). HTSeq--a Python framework to work with high-throughput sequencing data. Bioinformatics 31, 166–169. 10.1093/bioinformatics/btu638.

64. Trapnell C, Williams B A, Pertea G, Mortazavi A, Kwan G, van Baren M J, Salzberg S L, Wold B J, and Pachter L (2010). Transcript assembly and quantification by RNA-Seq reveals unannotated transcripts and isoform switching during cell differentiation. Nat. Biotechnol. 28, 511–515. 10.1038/nbt.1621.

65. Koene J M, Jansen R F, Ter Maat A, and Chase R (2000). A conserved location for the central nervous system control of mating behaviour in gastropod molluscs: evidence from a terrestrial snail. Journal of Experimental Biology 203, 1071–1080.

66. Kanehisa M, Araki M, Goto S, Hattori M, Hirakawa M, Itoh M, Katayama T, Kawashima S, Okuda S, Tokimatsu T, et al. (2008). KEGG for linking genomes to life and the environment. Nucleic Acids Res. 36, D480–484. 10.1093/nar/gkm882.

