## Supplementary Information for "A Chitin-Binding Protein Ultra-highly expressed in the Outer Fold of Mantle is Related to Shell Colour in Pacific Oyster *Crassostrea gigas*"

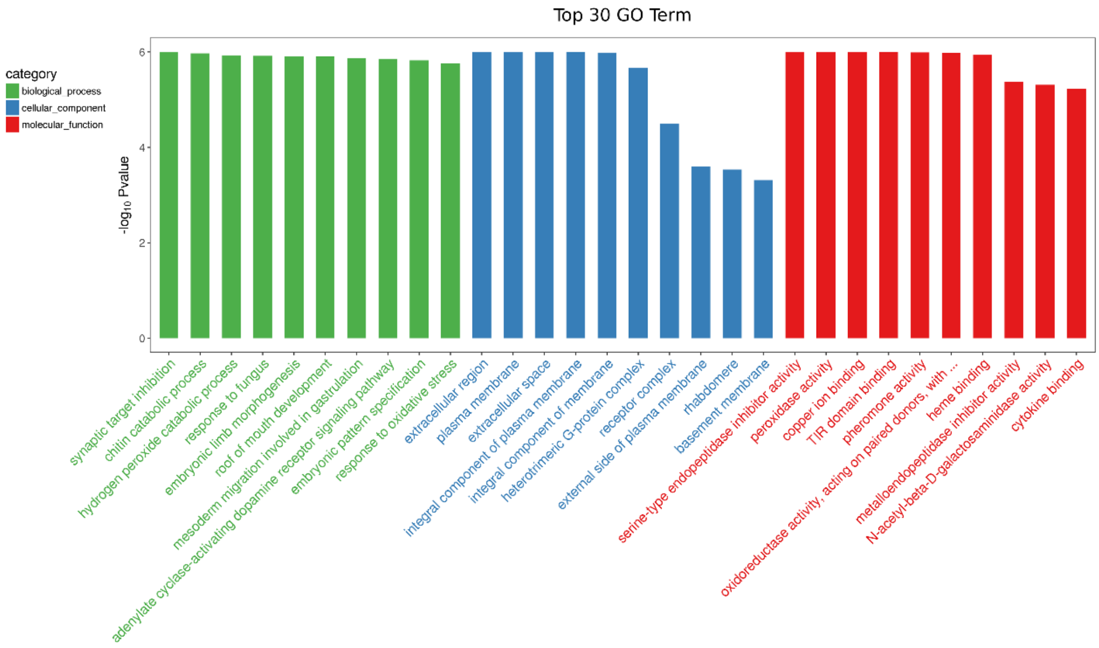


**Fig 1. GO enrichment for DEGs high expression in outer fold of mantle.**

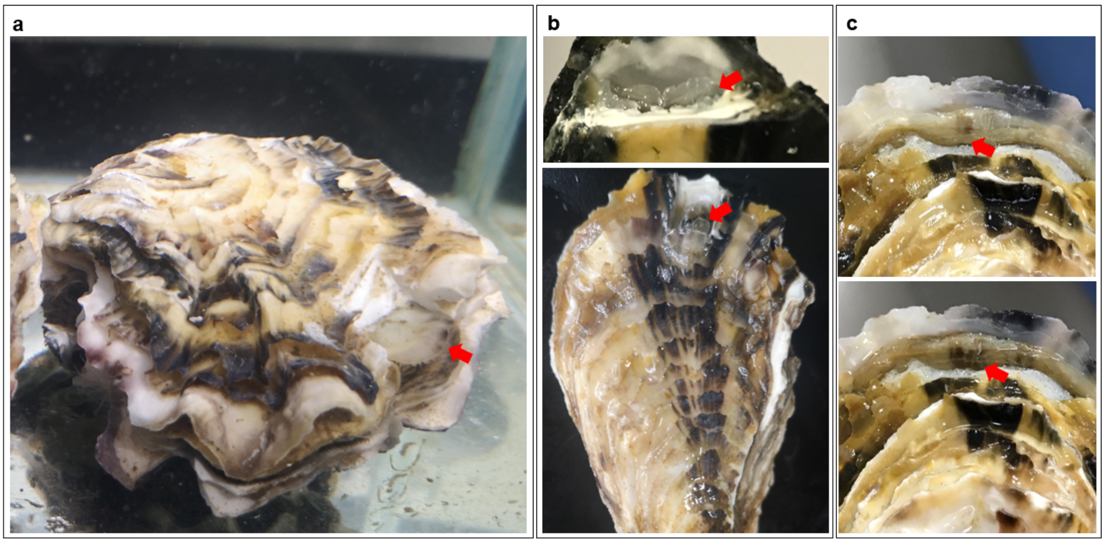


**Fig 2. Regeneration periostraca and pigmentation.** a: the mantle stretched out of the notching shell, closely attached to the regeneration periostraca; b: 0-36 h, regeneration periostraca, thin, transparent; c: 36-72 h, regeneration periostraca, thickened, pigmentation.

**Table 1. List of primers used for RT-qPCR analysis**

| Gene name | Primer sequences (5'–3') | Amplicon length (bp) | Accession No. |
|---|---|---|---|
| *CgCBP* | F: ATGCGAGCACTTCAGCACCAC | 140 | XM_011441463.3 |
| | R: CGTCTCCGAAGCCATCACAGC | | |
| *CgRS18* | F: GCCATCAAGGGTATCGGTAGAC | 168 | NM_001305367.1 |
| | R: CTGCCTGTTAAGGAACCAGTCAG | | |

**Table 2. Primers used to construct protein-expressing plasmids**

| Gene name | Primer sequences (5'–3') | Accession No. |
|---|---|---|
| CgCBP-myc | F: CATGGAGGCCCGAATTGGTATTTGTGAGGGATTATTTGG | XM_011441463.3 |
| | R: CTCGGTCGACCGAATTTTACATGTCCATTTCTTTAAAATCT | |
| CgNC1-myc | F: CATGGAGGCCCGAATTATGAGTGAGACAGCATCCCAAACG | LOC105338795 |
| | R: CTCGGTCGACCGAATTTCATTTCTGACTAGGTCCGCCAGC | |
| CgNC2-myc | F: CATGGAGGCCCGAATTATGTCGTCCACAAATACAAAGCC | LOC105347124 |
| | R: CTCGGTCGACCGAATTTCACTGTGGCTTGTAATTCACC | |

**Table 3. Sequences of dsRNA**

| dsRNA name | Sequences (5'–3') | Species |
|---|---|---|
| CgCBP | F: CCAGAGAGCUGCGCUAAAUTT | *Crassostrea gigas* |
| | R: AUUUAGCGCAGCUCUCUGGTT | |
| NC | F: UUCUCCGAACGUGUCACGUTT | *Caenorhabditis elegans* |
| | R: ACGUGACACGUUCGGAGAATT | |

**Table 4. GO enrichment analysis of high-expression proteins in black periostraca**

| GO_id | Term | Category | Pval | Protein_Gene |
|---|---|---|---|---|
| GO:0042438 | melanin biosynthetic process | BP | 2.47E-13 | K1RFL7, K1QAP4, K1PS92, K1PI66, K1R932, K1QE57, K1RLR4 |
| GO:0033162 | melanosome membrane | CC | 2.47E-13 | K1RFL7, K1QAP4, K1PS92, K1PI66, K1R932, K1QE57, K1RLR4 |
| GO:0016716 | oxidoreductase activity, acting on paired donors, with incorporation or reduction of molecular oxygen, another compound as one donor, and incorporation of one atom of oxygen | MF | 2.47E-13 | K1RFL7, K1QAP4, K1PS92, K1PI66, K1R932, K1QE57, K1RLR4 |
| GO:0008061 | chitin binding | MF | 5.85E-12 | K1RC30, K1R5I3, K1RC35, K1PPV2, K1QJK2, K1R3V2, K1RN92, K1PHQ7, K1RVI4 |
| GO:0005507 | copper ion binding | MF | 3.29E-10 | K1RFL7, K1QAP4, K1PS92, K1QQA2, K1PI66, K1R932, K1QE57, K1RLR4 |
| GO:0005975 | carbohydrate metabolic process | BP | 5.70E-06 | K1RC30, K1R5I3, K1QQL7, K1RC35, K1PHM5, K1RVI4 |
| GO:0004601 | peroxidase activity | MF | 9.24E-06 | K1QLH8, K1Q5E5, K1QCW5, K1Q633 |
| GO:0006979 | response to oxidative stress | BP | 1.76E-05 | |
| GO:0005576 | extracellular region | CC | 4.07E-05 | K1PR35, K1RC35, K1PPV2, K1QJK2, K1R3V2, K1RN92, K1PHQ7 |
| GO:0020037 | heme binding | MF | 0.000517 | K1QLH8, K1RFW1, K1Q5E5, K1QCW5, K1Q633 |
| GO:0003756 | protein disulfide isomerase activity | MF | 0.000229 | K1Q7T5, K1Q6X5 |
| GO:0005788 | endoplasmic reticulum lumen | CC | 0.000447 | K1Q7T5, K1Q6X5 |
| GO:0016614 | oxidoreductase activity, acting on CH-OH group of donors | MF | 0.001369 | K1QZ34, K1QG43 |
| GO:0050660 | flavin adenine dinucleotide binding | MF | 0.017136 | K1QZ34, K1QG43 |
